# PolyA: a tool for adjudicating competing annotations of biological sequences

**DOI:** 10.1101/2021.02.13.430877

**Authors:** Kaitlin M. Carey, Robert Hubley, George T. Lesica, Daniel Olson, Jack W. Roddy, Jeb Rosen, Audrey Shingleton, Arian F. Smit, Travis J. Wheeler

## Abstract

Annotation of a biological sequence is usually performed by aligning that sequence to a database of known sequence elements. When that database contains elements that are highly similar to each other, the proper annotation may be ambiguous, because several entries in the database produce high-scoring alignments. Typical annotation methods work by assigning a label based on the candidate annotation with the highest alignment score; this can overstate annotation certainty, mislabel boundaries, and fails to identify large scale rearrangements or insertions within the annotated sequence. Here, we present a new software tool, PolyA, that adjudicates between competing alignment-based annotations by computing estimates of annotation confidence, identifying a trace with maximal confidence, and recursively splicing/stitching inserted elements. PolyA communicates annotation certainty, identifies large scale rearrangements, and detects boundaries between neighboring elements.

## Introduction

Biological sequence annotation is the process of assigning labels to a sequence of nucleotides or amino acids, and is typically based on comparison (alignment) of such sequences to a database of known sequence elements. When the database contains elements that are similar to each other, many related elements may align with a high score to the same region of the target sequence being annotated, sometimes overlapping in non-trivial ways. In the face of competing annotations, software must ‘decide’ which is the true annotation at that location; we call this process *adjudication*. Current methods typically adjudicate the target as belonging to the element with the highest alignment score. This winner-takes-all approach overstates annotation certainty, struggles to determine the boundaries between neighboring partial element matches, and provides no basis for recognizing element nesting or rearrangements.

The existence of multiple related elements is common, even in databases designed to accumulate sequence instances into families. For example, in the Dfam [1] database of transposable element families, some abundant families are divided into highly similar subfamilies in order to represent family history or improve annotation sensitivity. Similarly, the Pfam [2] database of protein domains groups similar families into ’clans’, as does the Rfam [3] database of non-coding RNA families; in both cases, clan competition selects the longest and highest scoring hit when multiple clan members match the annotated sequence. For the purposes of exploring the value of confidence-based annotation adjudication, we focus on annotation of transposable elements (TEs) with RepeatMasker [4] and Repbase [5], a consensus sequence database of TE families with subfamily relationships similar to those of Dfam. To our knowledge, RepeatMasker (specifically its ProcessRepeats function) represents the most complex adjudication engine in use today, with expert knowledge built into the software to select between overlapping competing annotations, and even control the order of candidate alignments (when known) in order to recursively splice out inserted elements. In addition, RepeatMasker identifies cases in which multiple candidate annotations have nearly-identical scores, and reduces annotation specificity in some cases of ambiguity (e.g. labeling a sequence simply ‘Alu’ when several competing Alu subfamilies all share near-identical scores). Our goal in developing the software described here has been to replace this domain-specific framework with one that works with generic alignment tools and scoring matrices, requiring no expert knowledge of the underlying family structure, and improves representations of confidence, overlap, recombination, and insertions/stitching.

We have previously [6] described a simple method for computing confidence in (sub)family membership in the face of competing sequence alignments, making it possible to report confidence values for sequence labels, rather than simply reporting a single ‘winner’ from all possible matches. Here we extend this confidence scoring method to compute position-specific annotation based on the ensemble of competing alignments, assigning labels (with annotation confidence) to each position of the target sequence. Though our method is intended to be generally applicable to amino acid and nucleotide annotation, we present it in the context of the challenging domain of transposable element annotation adjudication. We call our software PolyA, short for ‘AAAAAAAAAAAAAAAA: Automatically Adjudicate Any And All Arbitrary Annotations, Astutely Adjoin Abutting Alignments, And Also Amputate Anything Amiss’. To our knowledge, PolyA is the first tool that computes a measure of annotation confidence in the face of competing alignments, and the first to establish a general framework for selecting the boundary cutoffs between overlapping annotations.

Our analysis focuses on transposable elements because these present all of the annotation challenges that PolyA is intended to resolve. Some families of TEs are represented by multiple closely-related subfamilies, which typically represent the family’s replication history (though are occasionally designed simply to improve annotation coverage). High levels of sequence similarity between subfamilies can cause several database sequences to align well to a specific genomic sequence, necessitating adjudication between competing annotations. An addition, integration of one element into another is a hallmark of TE activity, so that split sequences and unclear boundaries are very common. Furthermore, due to sequence similarity between elements, rearrangements and gene conversion are common in TEs. Current TE annotation is often performed using a tool called RepeatMasker (RM) [4], which uses the alignment score winner-takes-all approach when annotating fragments, then employs a complex expert system to stitch together the elements fragmented by other TE insertions.

In the following sections, we introduce PolyA, and demonstrate that it (i) reports a meaningful measure of annotation certainty, (ii) recognizes many instances of homologous recombination, (iii) effectively stitches sequence element segments resulting from nested insertions, (iv) accurately locates boundaries between overlapping candidate annotations, and (v) yields adjudicated annotation results that agree with those of RepeatMasker without resorting to complex and domain-specific expert system logic. We follow with a complete description of the methods supporting the observed results. PolyA is available for download at https://github.com/TravisWheelerLab/PolyA.

## Results

To annotate with PolyA, the target (to-be-annotated) sequence should be aligned to all the sequences in the annotation database. Alignment may be performed with any sequence-to-sequence alignment tool that depends on a scoring matrix (e.g. blast [7] or cross_match [8]). For the results presented here, this alignment is performed with cross_match, since this is the most sensitive alignment method used in RepeatMasker for consensus sequences; the annotation database is the Repbase RepeatMasker Edition library (a future version of PolyA will accept alignments with profile hidden Markov model databases such as Dfam and Pfam; see Discussion). PolyA takes as input (i) the collection of alignments between the target sequences and the database, and (ii) a reference to the scoring matrix and gap parameters used to produce each alignment. PolyA computes a measure of the confidence with which each position of the target can be assigned to each competing annotation candidate (see Methods). These position-specific confidence estimates for all competing annotations support identification of transition points generated by forces such as recombination and transposable element integration, and enable boundary detection between adjacent elements with overlapping competing annotations. A simple dynamic programming approach identifies a highest-confidence path through competing annotations, assigning labels to each position of the target sequence. This is followed by a stage that iteratively identifies insertion events and stitches the segments that were split by such insertions. The result is an annotation adjudication that correctly addresses events that typically cause incorrect labeling, and also discloses a measure of annotation certainty. See Methods for details.

To evaluate the efficacy of PolyA, we constructed a variety of artificial sequences with known origin, insertion activity, and recombination breakpoints. In each case, artificial sequences were constructed by mutating consensus sequences from Repbase, according to average substitution, insertion, and deletion rates acquired from the RepeatMasker hg38.fa.out annotation file [4], then inserting, trimming, and recombining as appropriate for the specific scenario being assessed. To explore annotation of true genomic sequence, we also compared PolyA-adjudicated annotation of the human genome with the annotation produced by RepeatMasker, through both summary statistics and manual examination of numerous complex genomics regions.

### Annotation Confidence

In [6] we introduced a simple mechanism for computing annotation confidence when several competing candidates produce alignments to a sequence window (see Methods). This confidence measure decreases as mutational load of an annotated sequence increases. To demonstrate this, we selected the AluY subfamily consensus sequence, and for each integer value in the range *p* ∈ [1‥50], generated 100 mutated copies of AluY in which *p* percent of nucleotides were randomly selected, and modified (uniformly) to a different nucleotide. Each mutated instance was aligned to all Repbase Alu subfamily consensus sequences using cross_match, and Eq 2 was applied to compute the confidence with which the instance was assigned an annotation of AluY. Fig 1 provides the average confidence for each bin of 100 (dark line), along with a shaded region indicating an interval of 1 standard deviation. This plot provides a sense of the decay in confidence expected as sequence divergence increases; specific confidence details for a particular (sub)family will depend on the number and relationship of similar subfamilies.

**Figure 1.**
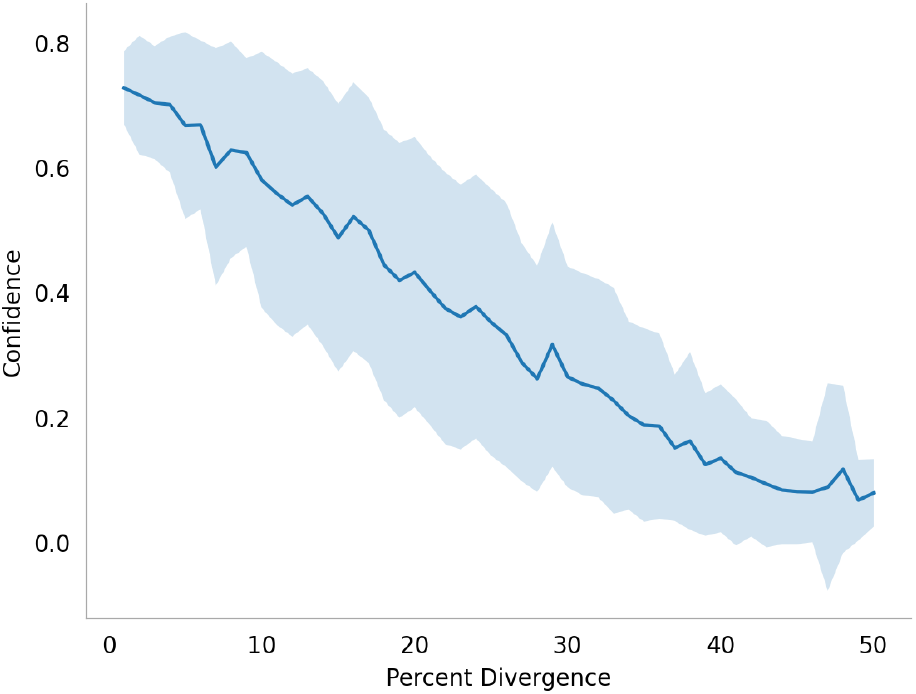
Confidence as a function of sequence divergence. Beginning with the Repbase consensus sequence for the AluJr subfamily, mutated instances were created with a range of percent substitutions, with 100 copies for each percentage bin. Each mutated instance was aligned to all Alu subfamilies using cross_match (25p41g matrix, gap_init=−25, and gap_ext=−5). For each sequence, the confidence in AluJr as the correct annotation was captured. The dark blue line shows the average AluJr confidence per bin, and the shaded region shows the range of a single standard deviation.

### Selecting Change-point for Overlapping Annotation Candidates

When faced with competing, overlapping annotation candidates, PolyA infers a precise intermediate boundary between overlapping annotation candidates by computing per-position confidence values and identifying a highest-confidence trace. This approach is effective for recombinations and insertions as described below, as well as for overlapping neighbors as demonstrated in Fig 2, in which the tail of an L1M1_orf2 candidate overlaps with several competing Alu alignments. The bottom of Fig 2 represents the position-specific confidence values as a heatmap, with green representing high confidence and purple representing low confidence (Note: the Alu shows long stretches of low confidence because that region of the genome is equally-well explained by several other Alu subfamilies - all of them share equally-low confidence over that region).

**Figure 2.**
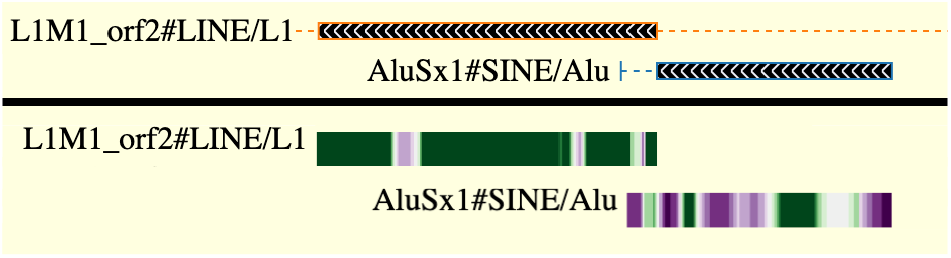
Detecting the change-point between overlapping elements; demonstration of confidence heatmap. This region of the human genome (hg38, chr10:58800-59500) contains a fragment of an L1M1 element, overlapping the tail of an AluSx1 candidate annotation. <u>The heatmap on the bottom</u> represents the position-specific confidence computed in PolyA, with dark green = high confidence, dark purple = low confidence, and moderate confidence represented by lighter colors and white. To limit visual clutter, this heatmap shows only the confidence heatmaps for the two ’winning’ annotations; others are hidden from view. <u>The top half of the figure</u> shows the final PolyA adjudication, in which the region in which the competing annotations overlap is assigned to the higher-confidence L1M1 annotation.

### Recombination

The presence of many highly-similar TE instances in a genome leads to common occurrence of non-allelic homologous recombination [9, 10], in which the initial sequence *a* is replaced in part by a subsequence of some homolog *b*. If *a* and *b* belong to different subfamilies *A* and *B*, respectively, then the sequence alignment step of annotation is likely to find near-full-length alignments of the sequence to database representatives of both *A* and *B*; the alignment to the *A* representative will be relatively poor in the region that was replaced by a stretch of sequence *b*, and vice versa. Using the standard adjudication process, the alignment with the higher score will be selected, and the existence of recombination will not be annotated.

PolyA computes position-specific annotation confidence estimates, and uses them to infer the presence, location, and identity of such recombinations. Fig 3 shows an example of an apparent recombination that is missed by the standard best-score-wins approach, but is detected by PolyA.

**Figure 3.**
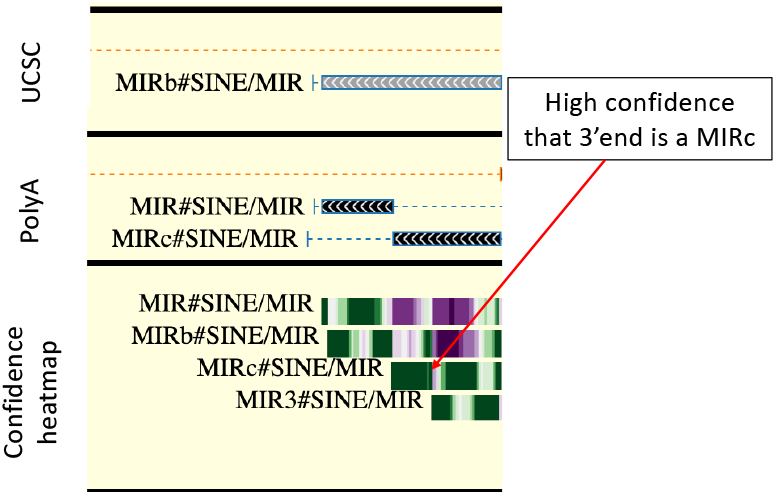
Recombination example. <u>The top panel</u> shows the RepeatMasker-adjudicated annotation of human (hg38) chr19:15304678-15304839 as a full-length MIRb element. <u>The bottom panel</u> presents a heatmap of the windowed average confidence that serves as the basis of PolyA adjudication, which highlights that in the 3’ end of the region, MIRc is the preferred annotation with high confidence. <u>The middle panel</u> shows that PolyA recognizes a recombination between MIR and MIRc at roughly the midpoint of the sequence window. The standard RepeatMasker preference for MIRb is due to the fact that it has the largest score of all candidates (MIRb=349, MIR=280, MIRc=290, MIR3=196). In the 3’ half of the annotated region beginning at position 15304742, MIRb’s score is 269, while MIRc’s score is 290.

Identification of homologous recombination is sure to be imperfect: (1) recombination events with highly similar donor and acceptor sequences, or with short donor segments, are relatively unlikely to be recognized, and (2) non-recombined sequence elements that are highly diverged from their appropriate consensus sequence are at increased risk of recombination false positive (due to reduced confidence as in Fig 1). We aimed to quantify sensitivity and false annotation by generating a large number of simulated sequences. In the following sections we describe the creation of these simulated benchmarks, and accompanying assessment. For all benchmarks based on simulated sequences, alignments were performed using cross_match, with fixed parameters consisting of the 25p41g matrix (available at https://github.com/Dfam-consortium/RepeatModeler), gap_init=−25, and gap _ext=−5.

#### False Positive Recombination

We first sought to test the frequency with which PolyA incorrectly identifies a sequence as being the result of a recombination, specifically: how often does PolyA claim that a sequence element is derived from two subfamilies, even though the actual sequence is derived from a single subfamily. We began with Alu sequences: randomly selecting one Alu type (AluS, AluJ, AluY), then one subfamily within the selected type, among those found in Repbase. The consensus sequence corresponding to the selected subfamily was mutated with subfamily-specific rates of substitution, insertion, and deletion. Substitution rates were approximated from existing repeatmasker (RM) annotations, modeling transitions as twice as likely as transversions. Indel rates and lengths were also chosen based on RM annotations: an overall length distribution of insertions and deletions (capped at length 7) was computed from all RM alignments, and used as the basis for randomly selecting indel lengths; for each benchmark instance, such indels were accumulated until reaching family-specific indel averages determined from RM annotations. This was repeated 10,000 times, yielding 10,000 sequences that contain no recombination and should be identified as such. These sequences were aligned to all Alu subfamily consensus sequences using cross_match, and adjudicated using PolyA. Of these <u>Alu</u> sequences, <u>99.8% were correctly identified as being derived from a single subfamily</u>. The same procedure was performed to produce 10,000 L1 instances, randomly selecting among all Repbase <u>L1</u> subfamilies. Of these, <u>97.8% were correctly identified as having a single source</u>.

#### True Positive Recombination

To assess PolyA’s sensitivity in detecting the aftermath of recombination, we performed experiments similar to those above, but now *with* recombination. Beginning with Alu, we: (i) selected a pair of distinct Alu subfamily consensus sequences *a* and *b*, with uniform probability as described above, (ii) aligned *a* and *b* using cross_match, and (iii) identified the middle position *i* of the alignment. If in the resulting sequence halves, either was shorter than 50 nucleotides, the simulation for this pair was aborted. With surviving simulations, (iv) both *a* and *b* were mutated as above, then (v) a recombinant sequence was created, made up of the prefix of *a* up to the nucleotide corresponding to position *i* of the alignment and the suffix of *b* beginning just after the *i*th position. This was repeated until 10,000 sequences were successfully generated, producing 10,000 simulated recombined sequences. We call these simulated sequences, with one end from *a* and one end from *b*, ‘single-recombinations’. Each resulting sequence was aligned to all Repbase Alu consensus sequences with cross_match, residue-specific confidence estimates were computed, and these were used to label the entire sequence allowing for recombination.

From these simulated <u>Alu single recombinants 55.1% were correctly identified</u> as recombinants. Among these, 65.2% were assigned to the correct subfamily on both ends of the recombination, while 96.9% were correctly assigned on at least one half.

A similar procedure was followed to produce L1 single-recombinations. Because Repbase L1 sequences are fragments of a full L1 (corresponding to 3’ end, 5’ end, and internal ORF region), many pairs do not meaningfully overlap. Furthermore, highly-divergent sequences are unlikely to produce homologous recombination. We performed an all-vs-all alignment of Repbase L1 consensus sequences, and identified 1553 alignment regions with length ≥ 100 bp on both sequences and ≥ 90% identity. With a given sequence-pair, we selected a random break-point and produced a recombinant as above, repeating 7 times for each pair, yielding a total of 10,871 L1 single-recombinations. These were each aligned using cross_match to the full contingent of Repbase L1 consensus sequences, with resulting alignments input to PolyA. Because PolyA adjudicates between annotation candidates produced by the alignment tool, it will fail to assign a correct family to a region if the input from the precursor alignment software does not include the correct family. This occurred in 5 L1 inputs. These have no possibility of being correctly adjudicated and were removed from the analysis. Of the remaining inputs, <u>94.8% of simulated L1 recombinants were correctly identified</u> as recombinants. Among these, 93.6% were assigned to the correct subfamily on both ends of the recombination, while 99.8% were correctly assigned on at least one half.

We repeated the above experiment for sequences simulating gene conversion, in which an internal segment of the original sequence is replaced by sequence from a related subfamily. Specifically, we identified two subfamilies and aligned their consensus sequences *a* and *b* as before, then *two* recombination points *i* and *j* were identified at positions 1/3 and 2/3 into the alignment, aborting construction of sequences where all segments are not atleast 50 nucleotides in length. For the remaining simulations both sequences were mutated, and the segment from sequence *a* corresponding to alignment positions *i* through *j* was replaced with the corresponding segment from sequence *b*. As before, 10,000 simulated sequences were produced from the Alu family, and for the L1 family 1228 alignment regions ≥ 90% identity and that fulfill the length requirement were identified. Repeating the gene conversion sequence construction described above 9 times for each pair we produced 11052 sequences. For these ‘internal recombinations’, 4 L1s were thrown out because input alignments from cross_match did not include the correct subfamily alignments. Among surviving instances, only 2.7% of simulated Alus were correctly identified as containing an internal recombination, while 18.1% were correctly identified as resulting from *some* recombination (i.e. were annotated as a single-recombinant). For L1 simulations, 56.5% were correctly identified as being the result of internal recombination, while 63.3% were correctly identified is being the result of *some* recombination(s) (i.e. at least one part of the recombination was recognized).

In both experiments, the low recombination sensitivity in Alus is not particularly surprising, since the nucleotides typically used to discriminate one of the subfamilies may not even have made it into the final recombinant sequence. Particularly for internal recombinations, the component sequences are all typically short enough that discriminating nucleotides are likely to be missing, leading to annotation ambiguity.

### Stitching Annotations Fragmented by Nested Insertions

It is common to find a single instance of a younger TE (sub)family inserted inside an instance of an older TE (sub)family. A goal of PolyA is to support the automatic stitching of the segments separated by such (possibly-nested) insertions. This is achieved by repeatedly identifying an inserted element, splicing it from between the sequence fragments that it separated, then repeating the labeling process on the resulting stitched sequences. This procedure is repeated until no apparent insertions are observed (see Methods for details). Fig 4 shows a genomic region with nested inserted elements. PolyA first identified a series of confidently-annotated segments, then spliced the inner-most element (AluJr4), stitching the remaining sequence around the excision. The resulting full-length MSTA1 was then spliced, so that the surrounding LTR40a instance could be stitched, and identified in full.

**Figure 4.**
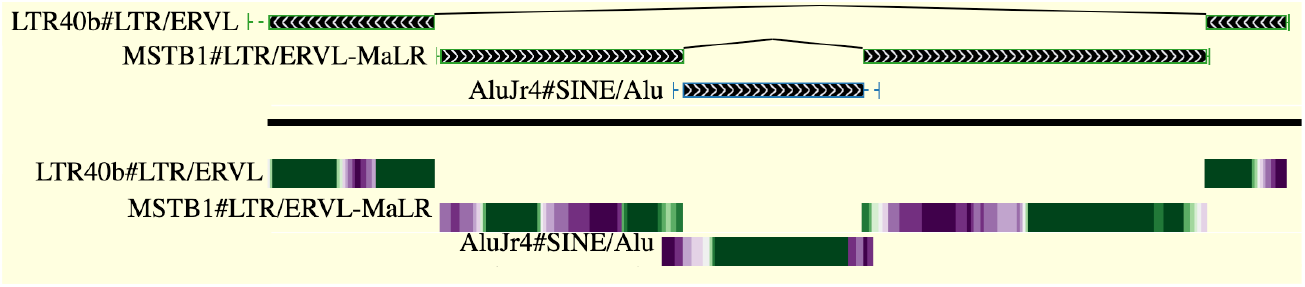
Example of nested insertions. This region of the human genome (hg38, chr11:11990878‥11991874) exemplifies nested transposable element insertions. Here, an instance of AluJr was inserted within an instance of MSTB1, which itself was inserted into an instance of LTR40b. The confidence heatmap is included for reference, and demonstrates a change-point decision in the context of nested annotations; heatmaps for competing annotations are not shown, in order to reduce visual clutter. PolyA automates the splicing of inserted elements and stitching of the surrounding split segments.

To assess the efficacy of PolyA’s annotation stitching mechanism, we simulated 3 classes of nested architectures, which we represent here as short strings: *ABA* (a single fragment of family *B* inserted into an instance of family *A*), *ABACA* (distinct fragments from families *B* and *C*,each inserted into an different location in an instance of family *A*), and *ABCBA* (a nested insertion, in which a fragment of family *C* is inserted into a fragment of family *B*, which itself is inserted into an instance of family *A*).

*ABA* sequences were simulated by creating 1,000 each of Alu1-Alu2-Alu1 (with Alu2 being a younger subfamily than Alu1), MER-Alu-MER, and MIR-MER-MIR amalgams. These are not meant to be exhaustive, but are representative of the kinds of insertions seen in the human genome. Alu families were selected as above, and others were selected randomly. In each case, the inner sequence was trimmed to retain the middle 2/3 of its length, and inserted into the middle of the outer sequence, splitting the outer sequence, but not replacing it. Trimming of the inner sequence was performed in order to replicate the common situation in TE annotation in which the boundary between inserted and surrounding sequence is unclear; this is a decidedly artificial arrangement, but it replicates the challenge faced during adjudication. Each of these 3,000 were aligned against the entire Repbase database, with results fed to PolyA. Of these, 182 sequences were filtered because the correct families were not among the cross_match results.

*ABACA* sequences were simulated by creating 1,000 each of LTR-L1^1^-LTR-L1^2^-LTR (in which L1^1^ and L1^2^ are two distinct L1 subfamilies), HAL1-Alu^1^-HAL1-Alu^2^-HAL1 (with distinct subfamilies Alu^1^ and Alu^2^), and L2-MER^1^-L2-MER^2^-L2 (with distinct subfamilies MER^1^ and MER^2^). In all other ways, these are produced as with the *ABA* format. Of these, 172 were filtered because the correct families were not among the cross_match results.

*ABCBA* sequences were simulated by creating 1,000 each of HAL1-MER-Alu-MER-HAL1, L4-MST-Alu-MST-L4, and L2-LTR-MST-LTR-L2. Of these, 122 were filtered due to failed cross_match results.

Table 1 shows that the correct nesting architecture was usually identified, as were the correct subfamilies.

**Table 1.** Accuracy results when annotating nested sequences.

|  | correct nesting architecture | correct families |
| --- | --- | --- |
| ABA | 93.3% | 83.4% |
| ABACA | 77.8% | 92.4% |
| ABCBA | 68.1% | 92.2% |

### Accurate Boundary Locations

To evaluate the accuracy of boundary point detection in PolyA, we computed the distance between estimated and true boundaries for all artificial sequences described in the experiments above (distance = |actual − detected|). The mean and median boundary detection error produced by PolyA is 5 and 9 nucleotides, respectively. Fig 5 shows the distribution of boundary detection error. Precise accuracy statistics depend on specifics of benchmark creation, alignment software choice, and aligner parameterization, but these results suggest that PolyA is effective at identifying cross-point boundaries.

**Figure 5.**
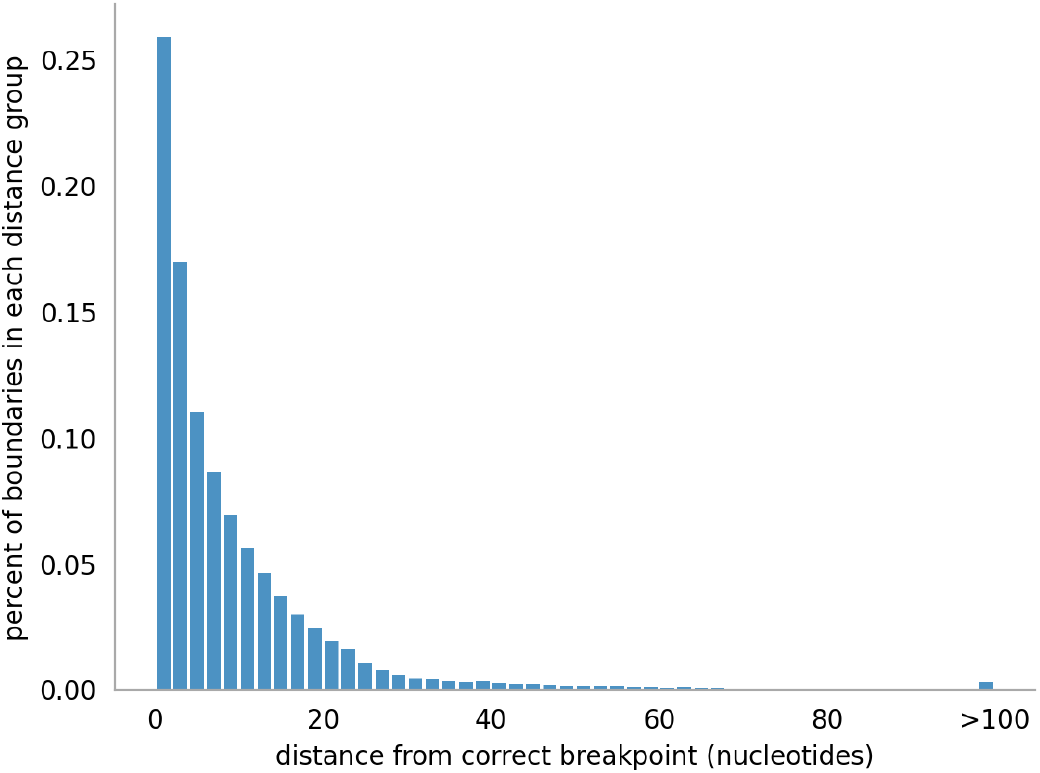
Boundary accuracy. To evaluate the accuracy of boundary point detection in PolyA, we computed the distance between estimated and true boundaries for all artificial sequences described above. All 24,961 sequences with correctly-labeled recombinations or nestings were used. The boundary error for each annotation was assigned to a bin of size 2. In 90% of all such annotations, the predicted boundary was within 20 nucleotides of the true boundary.

### Short tandem repeats

When annotating genomic sequences, it is common to mask short tandem repeats (STRs) prior to alignment-based annotation, either *hard-masking* (changing a masked region to a sequence of Ns so that alignment software assigns no score to alignments to the masked region) or *soft-masking* (marking a region such that that it will not serve as the seed of an alignment, but can be used in scoring after seeding is complete). PolyA accepts scored STR annotations from ULTRA [11] among candidate annotations. The left side of Fig 6 demonstrates how this may effectively allow an STR to out-compete a potential fragmentary family annotation (replacing a weak L1MC4 fragment annotation seen on UCSC based on RepeatMasker’s default adjudication) with the more appropriate STR call. The right side of Fig 6 shows an example of an STR candidate that is out-competed by the A-rich 3’ tail of an AluSz annotation.

**Figure 6.**
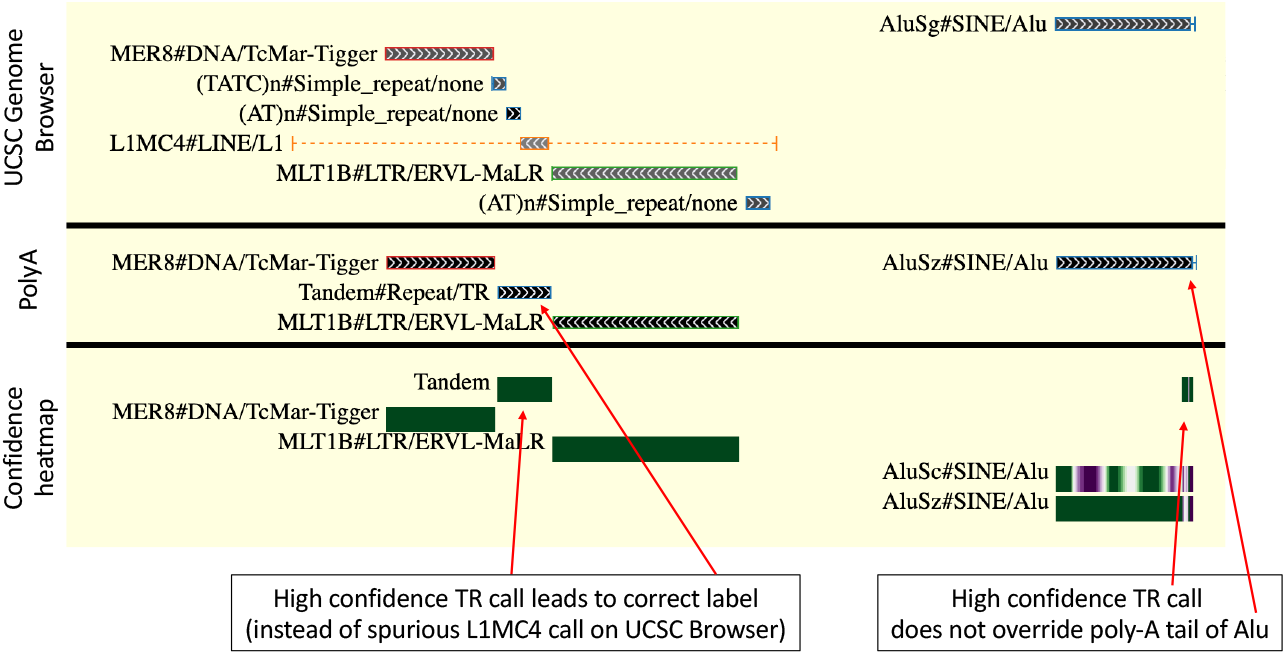
Adjudicating short tandem repeats. <u>The top section</u> shows the annotation of hg38, chr1:14632106‥146333918 as presented on the UCSC genome browser, which is derived from the standard adjudication results in RepeatMasker, using ProcessRepeats. This process aligns the library against genomic sequence that has first been masked for tandem repeats using TRF; it reports a short L1MC4 fragment of dubious accuracy (left), and a reasonable full-length AluSg annotation (right). <u>The middle section</u> shows the annotation produced when candidate annotations from RepeatMasker (TEs) and ULTRA (tandem repeats) are adjudicated using PolyA; it identifies the dubious L1MC4 fragment instead as a tandem repeat (left), and agrees on the Alu designation on the right (though subtle confidence differences lead to an AluSz assignment). <u>The bottom section</u> presents a confidence heatmap over the region. Of particular interest is the poly-A region on the far right of the plot; ULTRA correctly identifies this region as repetitive, but continuity with the preceding Alu annotation causes the region to be correctly labeled as part of the Alu.

In practice, it is likely still useful to soft-mask a genome prior to annotation, in order to limit excessive run time caused by evaluating alignments seeded in STR regions. Even so, by enabling direct competition between annotations from a library (by sequence alignment) or of STRs (as by ULTRA), PolyA provides substantially better fine-grained resolution of annotation near repetitive regions.

## Discussion

PolyA produces annotation confidence estimates for biological sequences based on an input of candidate sequence alignments and underlying scoring matrices. These confidence estimates are computed on a per-position basis, and used to infer transitions between overlapping annotations, including those caused by recombination and (possibly-nested) insertion of one element into another. We have demonstrated the efficacy of PolyA on multiple simulated scenarios, and shown that it produces reasonable results on the human genome. We are currently working to incorporate PolyA into the RepeatMasker software package for transposable element annotation.

PolyA often fails to identify recombination among instances of highly similar Alu elements - this is not surprising because (i) genomic Alu fragments are often short (e.g. less than 50 nucleotides), and thus unable to accumulate enough support to overcome transition penalties, and (ii) Alu subfamily sequences are highly similar, often differing by only a few nucleotides. We view this as a feature, not a failure. Importantly, PolyA produces confidence values for all annotations, and is therefore able to communicate the diminished confidence resulting from alignments to multiple highly-similar sequence elements, as with highly similar Alu subfamilies. We expect both the annotation tracing and confidence measures to contribute to future improvements in genome browser presentation of annotations.

In the current release, PolyA adjudicates annotations based on only nucleotide sequence-to-sequence alignments using scoring matrices, supplemented with tandem repeat annotations from ULTRA [11]. In a future release, PolyA will accept (i) annotations of protein sequences, and (ii) alignments made with profile hidden Markov models, which are the basis of sequence-family databases such as Dfam [1] and Pfam [2].

In its current form, PolyA naively adjudicates between candidate annotations without regard for additional information that expert systems currently use to improve annotation quality. In the future, we will explore approaches such as (i) adjusting transition penalties to prefer full length insertions when supported (akin to preferring global alignment [12]), and (ii) adjusting transition penalties based on relative apparent divergence of a sequence window to its candidate annotation.

## Methods

### Computing confidence in annotation of a full sequence

We can compute a measure of confidence that an annotated sequence belongs to a subfamily *i* by leveraging the probabilistic underpinnings of alignment scores. We have described these calculations in [6], but reproduce them here for completeness.

Given *Q* = *q*__1__*, q*__2__*, …, q__n__* competing subfamily annotations of genomic sequence *t*, we compute the confidence that *q__i__* is the correct label by normalizing over the probabilities of all competing labels:

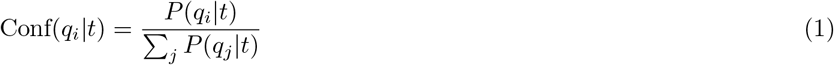

Assuming a uniform distribution over *Q*, *P* (*q__i__*|*t*) ∝ *P* (*t*|*q__i__*), so that

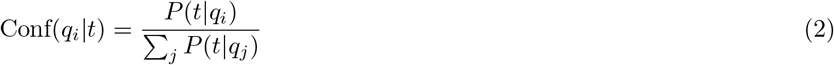

In scoring matrices used for sequence alignments, the score for aligning a pair of letters is a log odds ratio [13], typically scaled by factor *λ* then rounded to the nearest integer value. Under the simplifying assumption that gap costs map to probabilities [14], the overall alignment score corresponds to a scaled log of the ratio of the probability of observing *t* if it is homologous to *q__i__* vs the probability of observing *t* under a random (non-homology) model:

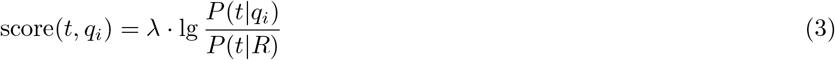

After straightforward algebraic manipulation

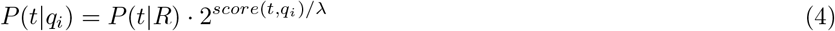

and following substitution into equation 2

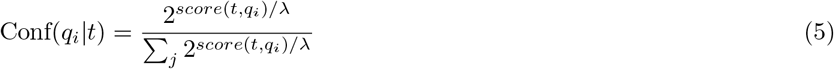

This derivation depends on the assumption that all competing family annotations are equally likely; in the case of non-uniform priors, with *P* (*q__k__*) defined as the prior probability of annotation *q__k__*, the computation may be adjusted as:

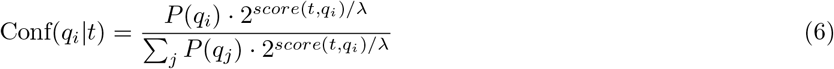

### Position-specific confidence, tracing through competing annotations

A natural mechanism for sub-family annotation would be to develop a jumping profile hidden Markov model (jpHMM) [15], representing the full contingent of candidate families, then to align genomic sequence to such a jpHMM. Posterior probabilities calculated on a nucleotide-resolution basis could work in lieu of the above calculations, and could also identify recombinations and boundary transitions between adjacent overlapping alignments. Unfortunately, such a model would be prohibitively computationally expensive. Here, we describe an efficient method for computing annotation confidence at position-level resolution, followed by a dynamic programming algorithm that traces a maximum confidence labeling of the annotated genome sequence.

To compute position-specific estimates of annotation confidence, we first modify the above full-sequence confidence calculations to compute confidence across length-*w* windows. For each position *j* in the annotated sequence of length *L*, we consider a window centered on *j* (trimmed at the ends) as follows: let *s* = max(1*, j* −⌊*w/*2⌋) and *e* = min(*L, j*+⌊*w/*2⌋), and compute the windowed confidence *Conf_i,j_* as above, considering only the scores of sub-alignments that cover sequence positions in the window (*s‥e*). By computing confidence within windows, our tool can account for the fact that one region of sequence is better explained by one candidate annotation, while another region is best explained by an alternate annotation.

The choice of window length is arbitrary - it should be long enough to smooth out the influence of individual mutations, but short enough to capture annotation transition points; PolyA sets the window length to *w* = 31 by default. In order to produce a more precise picture of annotation confidence, PolyA computes position-specific estimates for each position *j* by capturing an unweighted average of the confidence calculations of all windows that include position *j*:

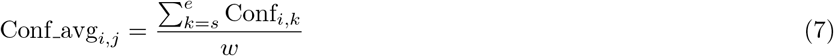

Based on these per-position confidence values, PolyA computes a maximum confidence trace through candidate annotations across the length of the annotated sequence, using the dynamic programming (DP) recurrence shown in Fig 7, in which each row is a candidate annotation *i*, and each column is a sequence position *j*.

**Figure 7.**
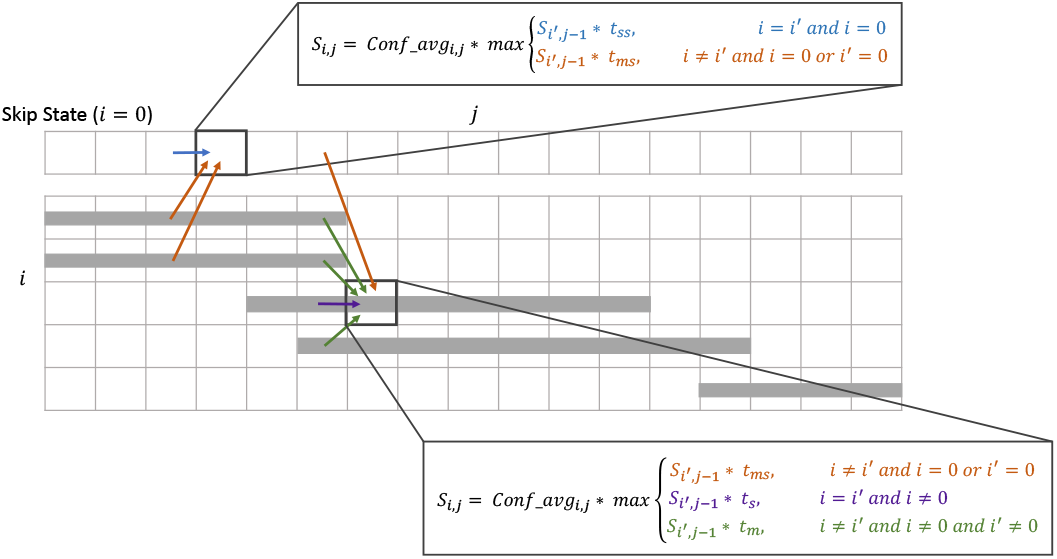
Dynamic programming recurrence. In the implicit dynamic-programming matrix, each column represents a position in the genome, and each row represents a candidate annotation family. In this figure, a gray bar corresponds to an alignment of the sequence for family i over a range of genome positions. Cells covered by a gray bar represent positions covered by an alignment - confidence (*Conf _avg__i,j__*) and score (*S__i,j__*) values are computed only these positions; other cells have an implicit value of zero. Because this describes a very sparse matrix, values are stored as a sparse set of (i,j,conf_avg) tuples. Transition probabilities are: *t_s_* (‘<u>s</u>tay in the same row’), *t_m_* (‘<u>m</u>ove to a new non-skip-state row’), *t_ms_* (‘<u>m</u>ove between the skip-state row and a non-ski state row’), and *t_ss_* (‘<u>s</u>tay in the skip-state row’). Default values are: *t_m_* = 1*e −* 55, 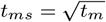, *t_s_* = 1 − *t_m_* − *t_ms_*, *t_ss_* = 1 − *t_ms_*.

Two additional candidate annotations are considered, beyond those provided in the form of sequence alignments. (1) Candidate short tandem repeats (STRs) are identified using the tool ULTRA [11], which provides per-position log-odds ratio scores that fit naturally into the PolyA confidence calculations; these are simply treated as an additional candidate annotation. (2) In some cases, a single low-quality annotation may extend slightly beyond a clearly-preferred annotation; if PolyA is forced to choose among available annotations, a small clip of this low-quality annotation will be selected. PolyA avoids this outcome by including what we call a *skip state*; when the annotation trace is in a skip state, the corresponding sequence is assigned no annotation. Skip states are given a special default score (such that they will out-compete low-quality candidate annotations).

To encourage annotation continuity, transitioning from one annotation candidate to a different annotation is heavily penalized. The penalty for transitioning to/from the skip state is reduced. Because an annotated sequence may exceed the length of any single candidate annotation, the Conf avg*i,j* matrix is sparse; the DP implementation performs correspondingly sparse calculations to avoid excessive time and space resource use.

A maximal trace through the sparse DP matrix described in Fig 7 yields an ordered series of labeled regions over the target sequence. These regions may identify transitions between adjacent elements and recombination or insertion events.

### Recursive splicing of insertions, and stitching of surrounding sequences

In the case that an annotated sequence contains an instance *B* of one family inserted into the middle of an instance *A* of another family, the above DP mechanism will annotate the sequence in a form *A*_1_*BA*_2_. PolyA takes steps to identify the relationship between *A*_1_ and *A*_2_ as ordered fragments of *A* (see Fig 8). To achieve this, a graph is created in which each labeled sequence block is represented as a vertex, and a directed edge is established between the vertices corresponding to adjacent segments. Additional edges are added to the graph for each pair of vertices corresponding to segments might share to the same family, specifically: (i) the alignments supporting the two segments are relatively co-linear in the label’s consensus sequence, and (ii) the label with highest confidence on one of the segments has some non-negligible (default ≥ 1%) confidence on the other segment (tested in both directions). In such a graph, an unbroken inserted element will appear as a vertex with in-degree and out-degree of 1. PolyA removes all such vertices from the graph, splices the corresponding positions from the confidence-based DP matrix, and repeats the DP trace. With the inserted sequence removed, the adjusted transition opportunities enable stitching of previously-separated segments. This process is iterated until no inserted sequences remain.

**Figure 8.**
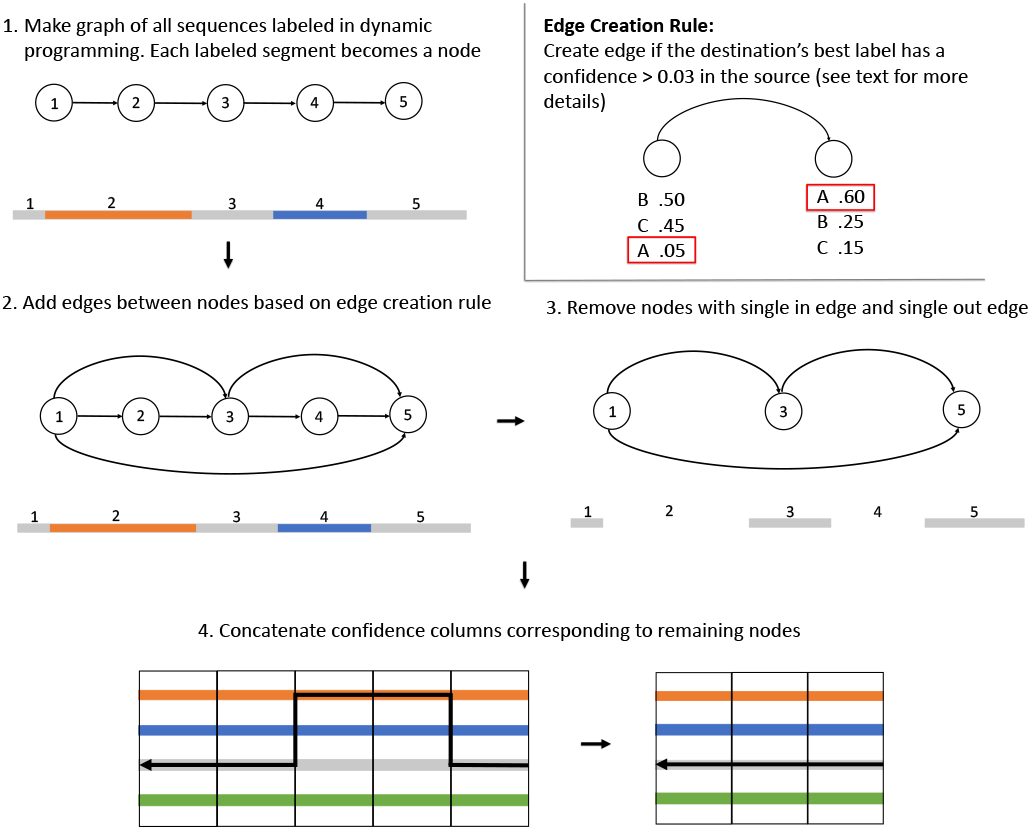
Splicing insertions. These images describe the simple graph algorithm used to identify inserted elements, splice them out, and stitch segments of the remaining sequence back together to be considered in another round of dynamic programming.

## Acknowledgements

We gratefully acknowledge the computational resources provided by the University of Montana’s Griz Shared Computing Cluster (GSCC), and the Montana Tech High Performance Computing Cluster.

## Abbreviations

TE: transposable element
STR: short tandem repeat
RM: RepeatMasker
DP: Dynamic programming

## Availability of data and materials

PolyA source code is available at https://github.com/TravisWheelerLab/PolyA

## Authors’ contributions

RH, AFS, JR, TJW established motivation for PolyA, and devised mechanisms for validation. TJW designed algorithms. KMC implemented the algorithm and performed analysis. KMC, GL, and AS were responsible for software development and release. AS and DO developed tandem repeat modifications and analysis. JWR produced software for visuals. JR devised the name. KMC and TJW led development of the manuscript, with contributions from others. All authors read and approved the final manuscript.

## Authors’ information

Department of Computer Science, University of Montana, Missoula MT, USA

Kaitlin M. Carey, George T. Lesica, Daniel Olson, Jack W. Roddy, Audrey Shingleton, Travis J. Wheeler

Institute for Systems Biology, Seattle WA, USA Robert Hubley, Jeb Rosen, Arian F. Smit

## References

1. Hubley, R., Finn, R.D., Clements, J., Eddy, S.R., Jones, T.A., Bao, W., Smit, A.F., Wheeler, T.J.: The Dfam database of repetitive dna families. Nucleic acids research 44(D1), 81–89 (2016)

2. El-Gebali, S., Mistry, J., Bateman, A., Eddy, S.R., Luciani, A., Potter, S.C., Qureshi, M., Richardson, L.J., Salazar, G.A., Smart, A., et al.: The Pfam protein families database in 2019. Nucleic acids research 47(D1), 427–432 (2019)

3. Nawrocki, E.P., Burge, S.W., Bateman, A., Daub, J., Eberhardt, R.Y., Eddy, S.R., Floden, E.W., Gardner, P.P., Jones, T.A., Tate, J., et al.: Rfam 12.0: updates to the rna families database. Nucleic acids research 43(D1), 130–137 (2015)

4. Smit, AFA. Hubley, R., Green, P.: RepeatMasker Open-4.0.2013-2015 (2013). http://www.repeatmasker.org/

5. Bao, W., Kojima, K.K., Kohany, O.: Repbase update, a database of repetitive elements in eukaryotic genomes. Mobile Dna 6(1), 11 (2015)

6. Carey, K.M., Patterson, G., Wheeler, T.J.: Transposable element subfamily annotation has a reproducibility problem. Mobile DNA 12(1), 1–9 (2021)

7. Altschul, S.F., Gish, W., Miller, W., Myers, E.W., Lipman, D.J.: Basic local alignment search tool. Journal of molecular biology 215(3), 403–410 (1990)

8. Green, P.: Phrap and cross_match. http://www.phrap.org/phredphrap/phrap.html

9. Fawcett, J.A., Innan, H.: The role of gene conversion between transposable elements in rewiring regulatory networks. Genome biology and evolution 11(7), 1723–1729 (2019)

10. Sung, P., Klein, H.: Mechanism of homologous recombination: mediators and helicases take on regulatory functions. Nature reviews Molecular cell biology 7(10), 739–750 (2006)

11. Olson, D., Wheeler, T.: Ultra: A model based tool to detect tandem repeats. In: Proceedings of the 2018 ACM International Conference on Bioinformatics, Computational Biology, and Health Informatics, pp. 37–46 (2018)

12. Brudno, M., Malde, S., Poliakov, A., Do, C.B., Couronne, O., Dubchak, I., Batzoglou, S.: Glocal alignment: finding rearrangements during alignment. Bioinformatics 19(suppl 1), 54–62 (2003)

13. Yu, Y.-K., Altschul, S.F.: The construction of amino acid substitution matrices for the comparison of proteins with non-standard compositions. Bioinformatics 21(7), 902–911 (2005)

14. Frith, M.C.: How sequence alignment scores correspond to probability models. Bioinformatics 36(2), 408–415 (2020)

15. Schultz, A.-K., Zhang, M., Leitner, T., Kuiken, C., Korber, B., Morgenstern, B., Stanke, M.: A jumping profile hidden markov model and applications to recombination sites in HIV and HCV genomes. BMC bioinformatics 7(1), 265 (2006)

